# Activating A1 adenosine receptor signaling boosts early pulmonary neutrophil recruitment in aged mice in response to *Streptococcus pneumoniae* infection

**DOI:** 10.1101/2024.01.08.574741

**Authors:** Shaunna R. Simmons, Sydney E. Herring, Essi Y.I Tchalla, Alexsandra P. Lenhard, Manmeet Bhalla, Elsa N. Bou Ghanem

## Abstract

**Background:** *Streptococcus pneumoniae* (pneumococcus) is a leading cause of pneumonia in older adults. Successful control of pneumococci requires robust pulmonary neutrophil influx early in infection. However, aging is associated with aberrant neutrophil recruitment and the mechanisms behind that are not understood. Here we explored how neutrophil recruitment following pneumococcal infection changes with age and the host pathways regulating this.

**Results:** Following pneumococcal infection there was a significant delay in early neutrophil recruitment to the lungs of aged mice. Neutrophils from aged mice showed defects in trans-endothelial migration *in vitro* compared to young controls. To understand the pathways involved, we examined immune modulatory extracellular adenosine (EAD) signaling, that is activated upon cellular damage. Signaling through the lower affinity A2A and A2B adenosine receptors had no effect on neutrophil recruitment to infected lungs. In contrast, inhibition of the high affinity A1 receptor in young mice blunted neutrophil recruitment to the lungs following infection. A1 receptor inhibition decreased expression of CXCR2 on circulating neutrophils, which is required for transendothelial migration. Indeed, A1 receptor signaling on neutrophils was required for their ability to migrate across endothelial cells in response to infection. Aging was not associated with defects in EAD production or receptor expression on neutrophils. However, agonism of A1 receptor in aged mice rescued the early defect in neutrophil migration to the lungs and improved control of bacterial burden.

**Conclusions:** This study suggests age-driven defects in EAD damage signaling can be targeted to rescue the delay in pulmonary neutrophil migration in response to bacterial pneumonia.

## BACKGROUND

*Streptococcus pneumoniae* (pneumococcus) are encapsulated, Gram positive bacteria that are responsible for an estimated 1.6 million deaths each year (1). *S. pneumoniae* remain the leading cause of community acquired bacterial pneumonia in individuals over the age of 65 (2–4). Disease and mortality caused by these bacteria persist despite available vaccines and antibiotics (3, 5, 6). Older adults are at an increased risk of infection and serious disease from pneumococcal pneumonia due to progressive deleterious molecular, cellular and systemic changes termed the “Hallmarks of Aging” (7, 8), that can affect the ability of the host to mount an efficient and balanced immune response. These hallmarks include aberrant genetic changes, altered metabolic responses, cellular senescence and chronic inflammation(7, 8). Cellular senescence affects several cell types including immune cells, causing them to be less efficient at pathogen clearance (7, 8). Cellular senescence is also associated with increased production of pro-inflammatory cytokines leading to an increase in basal inflammation known as inflammaging (7, 8). Increased inflammation with age contributes to a reduction in the ability of immune cells to respond to acute stimuli, further dysregulating the immune response (7–9). A better understanding of how immune cell responses change with host aging upon pneumococcal infection, and the signaling pathways that control these responses, is necessary to promote better protection against disease.

An important mediator of host protection following pneumococcal pneumonia is the polymorphonuclear leukocyte (PMNs) response. PMNs are the first cell to be recruited to the lungs following pneumococcal infection (10, 11) and this response is necessary for host protection. Individuals with neutropenia are more susceptible to pneumonia and experimental depletion of PMNs early in infection result in reduced ability to control bacterial numbers and accelerated death in murine models of infection (12, 13). Upon pulmonary infection, PMNs are recruited from the circulation into the lung tissue where they kill bacteria via several antimicrobial effector functions, such as release of preformed antimicrobial granules, reactive oxygen species, phagocytosis, and neutrophil extracellular traps (14). However, an effective PMN response in the pulmonary environment must be balanced, where PMNs need to be recruited in quickly to clear bacteria and then resolve. An exacerbated PMN response without resolution is damaging to host tissue due to the antimicrobial products produced by PMNs, and without proper resolution PMNs contribute to lung damage which worsens disease outcomes (12, 15). There is evidence that PMN influx to the lungs later in infection is dysregulated with age as biopsy samples from older adults show an increase in the percentage of PMNs in the lungs compared to younger patients following pneumococcal pneumonia, and this can be recapitulated in mouse models (16, 17). However, how neutrophil recruitment changes with age early in infection, when PMNs are beneficial to the host in clearing the pathogen, and the pathways controlling PMN influx remain unexplored.

We previously found that the production of extracellular adenosine (EAD) is crucial for regulating PMN influx to the lungs in young mice (10). Adenosine accumulates in the extracellular space as a result of cellular damage. Upon tissue injury by a variety of insults, including infection, ATP released from cells is metabolized to EAD by the sequential action of the extracellular ectonucleotidases CD39 and CD73 (18). At baseline, EAD levels are very low/ undetectable in extracellular spaces but can increase more than 10-fold upon infection (19). EAD can signal via 4 G-protein coupled receptors, A1, A2A, A2B and A3 that respond to EAD in a dose-dependent manner (18, 20). A1 is high affinity and is activated by lower levels of adenosine while A2A and A2B are lower affinity and require higher levels of adenosine to become activated (18). The different adenosine receptors are also coupled to different G proteins that can inhibit (Gi) or stimulate (Gs) adenylyl cyclase or activate phospholipase C (Gq), and therefore can have opposite effects on downstream signaling and cell function. These receptors are ubiquitously expressed on many cell types including PMNs (21).

Changes in EAD production and signaling occur with aging (22–27). Although the role of this pathway in immunosenescence remains largely unexplored, we previously found that aging is accompanied by changes in expression of EAD enzymes on murine PMNs at baseline and that EAD signaling controlled PMN antibacterial function (21, 28, 29). However, how the levels of extracellular adenosine and the receptors signaled through change after pneumococcal infection and across host age, and their role in PMN migration, is unknown. In this study, we asked how early PMN recruitment changes with host aging and if the extracellular adenosine pathway regulates that during bacterial pneumonia. We tested the hypothesis that signaling through the high affinity A1 adenosine receptor would be important early in pneumococcal infection when adenosine levels are lower to help draw the PMNs to the lung, while signaling through the lower affinity receptors may be required late in infection to help resolve PMN influx and that these dynamics change during host aging.

## METHODS

### Mice

Young (8-12 weeks) and old (18-22 months) C57BL/6 (B6) male mice were obtained from the National Institute on Aging colony or from Jackson Laboratories (Bar Harbor, ME). A2BR^-/-^ mice on a B6 background (B6.129P2-Adora2btm1Till/J) and A2AR^-/-^ mice on a Balb/c background (C;129S-Adora2atm1Jfc/J) were purchased from Jackson Laboratory and bred at our facility. Wild type C57BL/6J and Balb/cJ mice were used as controls. Since males are more susceptible to pneumococcal infection compared to females (30) and due to the limited availability of aged cohorts, experiments were performed in male mice. All mice were housed at the University at Buffalo in specific pathogen free housing for at least 2 weeks before use in experiments. All experiments were conducted in accordance with the Institutional Animal Care and Use Committee guidelines.

### Bacteria

*Streptococcus pneumoniae* TIGR4 strain (serotype 4) was a kind gift from Andrew Camilli. Briefly, bacteria were grown to mid-exponential phase at 37⁰C at 5% CO_2_ in Todd Hewitt broth supplemented with 0.5% yeast extract and oxyrase as previously described (20).

### PMN Isolation

Bone marrow cells were collected from the femurs and tibias of uninfected mice by flushing with RPMI supplemented with 10% FBS and 2mM EDTA. Red blood cells were then lysed, and remaining cells were washed, and resuspended in PBS. PMNs were separated via density centrifugation using histopaque 1119 and 1077 (Sigma) as previously described (20, 21, 31, 32). PMNs were then resuspended at the desired concentration in Hank’s Balanced Salt Solution/0.1% gelatin with no Ca^2+^ or Mg^2+^ and kept on ice until use. The purity of PMNs was confirmed using flow cytometry and 85-90% of enriched cells were positive for Ly6G and CD11b.

### Mouse Infections and Treatments

Mice were infected intratracheally (i.t.) with 50ul of *S. pneumoniae* TIGR4 at the indicated concentrations. Mice were first anesthetized using isoflurane and the bacterial inoculum pipetted into the trachea with the tongue gently pulled out to ensure direct delivery into the lungs. Following infection, mice were euthanized at the indicated time points, blood was collected by portal vein snip to determine bacteremia. Mice were then perfused with PBS through the right ventricle and the lungs harvested. Following harvest, the lungs were minced into small pieces, mixed well and half of each lung sample was homogenized in sterile PBS to plate for CFU, and the remaining half was digested and used for flow cytometry analysis. Lung and blood samples were diluted in sterile PBS and plated on blood agar to enumerate CFU. For experiments where the effect of A1 receptor signaling was measured, the A1 receptor inhibitor, 8-Cyclopentyl-1,3-dipropylxanthine (DPCPX)(28, 33, 34) or A1 receptor agonist 2-Chloro-N6-cyclopentyladenosine (35) were purchased from Sigma Aldrich, dissolved in DMSO and filter sterilized by passing through a 0.22μm filter prior to use. Mice were given intraperitoneal (i.p.) injections of 1mg/kg of the A1 inhibitor or 0.1mg/kg of the A1 agonist or vehicle control at −1 and 0 (immediately before challenge) days relative to infection. For experiments where the effect of A2A receptor signaling was measured, the A2A receptor inhibitor, 3,7-Dimethyl-1-propargylxanthine (D134) (36), or A2A receptor agonist CGS21680 (37), were purchased from Sigma, dissolved in DMSO and filter sterilized by passing through a 0.22 μm filter prior to use. Mice were given i.p injections of 5mg/kg of the A2A antagonist or 2mg/kg of the A2A agonist or vehicle control at −18 and 0 hours, immediately before challenge for pre-treated (designated as PRE) and 18 hours post challenge for post-treated groups (designated as POST). Control animals were treated with vehicle control.

### Flow Cytometry

The lungs were digested into a single cell suspension in RPMI 1640 containing 10% FBS, 2mg/mL type 2 collagenase and 50 U/mL DNase (Worthington) at 37°C/ 5% CO_2_ as previously described (20). Blood was collected by portal vein snips in EDTA as an anticoagulant. Red blood cells were removed with a hypotonic lysis buffer (Lonza). Cells were resuspended in FACS buffer (HBSS/ 1% FBS), treated with Fc block (anti-mouse clone 2.4G2) and stained with the following antibodies purchased from BD Biosciences or Invitrogen: Anti-Ly6G (IA8), anti-CD11b (M1/70), Anti-CD45 (30-F11). For adenosine receptor staining, the BD Cytofix/Cytoperm kit was used to permeabilize the cells and the cells stained with the following unconjugated rabbit anti-adenosine receptor antibodies from Abcam as previously described (21): A2a (ab3461), A2b (ab222901), A3 (ab203298) and A1 (ab82477). Rabbit polyclonal IgG (ab37415) was used as an isotype control and secondary PE-conjugated anti-Rabbit IgG was used (12473981; Invitrogen). Fluorescent intensities were measured on a BD Fortessa and at least 20,000 events were analyzed using FloJo.

### Trans Endothelial Migration Assay

To determine trans-endothelial migration of PMNs *in vitro*, C57BL/6 murine primary lung endothelial cells were grown according to manufacturer’s protocol (Cell Biologics) and 5μm transwells (Corning) were seeded at 10^6^ cells/transwell and grown for 2 days at 37°C/ 5% CO_2_ until confluent. The bottom of the transwell was infected with *S. pneumoniae* TIGR4 at an MOI of 40. PMNs were isolated from the bone marrow of indicated mouse strains and 5×10^5^ PMNs placed in the top of the transwell and incubated at 37°C at 5% CO2 for 1.5 hours. The number of PMNs that migrated through the endothelial layer into the bottom of the transwell was determined by collecting the cells from the bottom well and lysing them with 10% Triton X-100 to release MPO. MPO was measured in each well by using 2,2′-Azino-bis(3-ethylbenzothiazoline-6-sulfonic acid) diammonium salt (ABST) in the presence of hydrogen peroxide (10) and serial dilutions of a known numbers of PMNs were used to establish a standard curve, which was then used to quantify the number of migrated neutrophils as previously described (10).

### Measurement of Adenosine

Mice were infected i.t with 2.5×10^4^ CFU *S. pneumoniae* TIGR4. Following infection, mice were euthanized at indicated time points and blood was collected and placed into microtainer tubes (BD) and centrifuged at 9000 rpm to collect sera. Circulating levels of adenosine were measured using the Adenosine Assay Kit (Fluorometric) from BioVision per manufacturer’s instructions. Results were read using a Biotek plate reader.

### Statistics

Statistics were analyzed using Prism 9 (GraphPad). Bar graph and scatter plot data are presented as Mean +/-SD, and line graph data are presented as Mean +/-SEM. All data were tested for normality using Shapiro-Wilk test prior to statistical analysis. Significant difference between groups was determined by Student’s t-test, Mann-Whitney, Kruskal Wallis or one-way ANOVA as indicated. Difference in survival was determined by the log-rank (Mantel-Cox) test. *p* values *<0.05* were considered significant. * indicates *p <0.05*, ** indicates *p <0.001*, *** indicates *p<0.0001*.

## RESULTS

### Rapid PMN influx to the lungs following *S. pneumoniae* infection is impaired with aging

To determine how PMN influx to the lungs changes with age throughout the course of pneumococcal infection, young and aged mice were infected i.t. with 2.5×10^4^ colony forming units (CFU) of *S. pneumoniae* TIGR4 and PMN recruitment to the lungs was measured by flow cytometry over time following infection. Young mice had significant PMN influx to the lungs within 6 hours of infection, and further increase in pulmonary PMN numbers by 18 and 48 hours post infection (hpi) (**Fig 1A**). With age there was a delay in PMN influx early in infection with no increase from baseline by 6 hours, but a significant increase by 18 and 48 hpi (**Fig 1A**). When compared to young controls, aged mice had approximately 4-fold lower PMNs in the lungs at 6 hpi (**Fig 1A**). By 18 and 48 hpi, PMNs were present in the lungs of aged mice at similar levels to young mice. This delay in PMN influx to the lungs with age was associated with higher pneumococcal CFU in the lungs throughout the course of infection (**Fig 1B**) and significantly higher bacteremia at 18 hpi (**Fig 1C**). When normalized to bacterial burden, aged mice displayed significantly reduced PMN numbers in the lungs compared to young controls at 6 hours and significantly higher numbers of PMNs at 48 hpi (**Fig S1A)**. These data indicate that with age there is an initial delay in neutrophil influx to the lungs following pneumococcal infection that is associated with impaired bacterial control.

**Figure 1:**
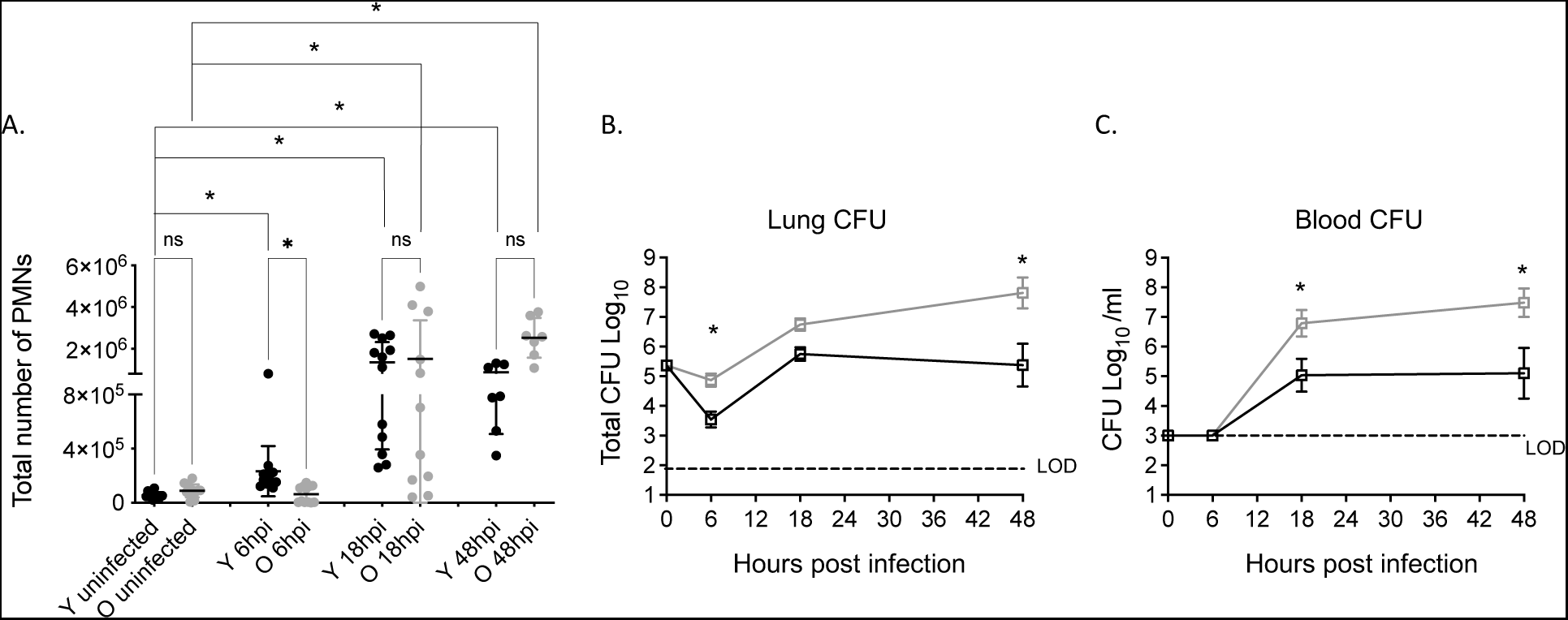
Old mice have delayed early pulmonary PMN influx. Young and old male C57BL/6 mice were infected i.t. with 5×10^4^ CFU *S. pneumoniae* TIGR4. (A) Lungs were harvested, digested, and stained for PMNs and analyzed by flow cytometry at the indicated timepoints following infection. (B) Lung and blood (C) were collected, and samples plated on blood agar and CFU was enumerated at the indicated timepoint post infection. Data are pooled from 6 separate experiments with a total of n=7 mice at 48-hour time point and n=12 mice at all other time points. Asterix indicates significant difference between indicated groups and as calculated by (A) Kruskal Wallis followed by Dunn’s multiple comparison test and (B and C) one-way ANOVA followed by Sidak’s multiple comparison test. ns indicates non-significant.

### PMN trans-endothelial migration in response to *S. pneumoniae* is impaired with aging

For PMNs to enter the lungs to control pneumococcal infection, these cells must be recruited from the circulation and migrate through the endothelium to the infected pulmonary tissue. To better understand the age-related impairment in PMN influx, we measured neutrophil trans-endothelial migration. Using an *in vitro* transwell model, we measured neutrophil migration across primary mouse lung endothelial cells (isolated from young mice) in response to *S. pneumoniae* as a stimulus. At baseline there was no difference in neutrophil migration, however upon infection, there was a significant decline in the number of PMNs that migrated through the endothelial cell layer when PMNs were isolated from aged mice compared to young controls (**Fig 2**). These data suggest that there is an intrinsic defect in trans-endothelial migration by PMNs from aged mice.

**Figure 2:**
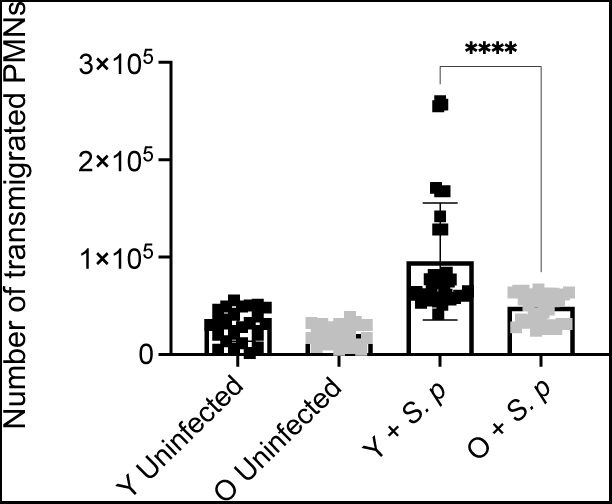
PMNs from old mice have a defect in trans-endothelial migration. PMNs were isolated from the bone marrow of young and old male C57BL/6 mice. Transwells were seeded with murine primary lung endothelial cells and the bottom of the transwell infected with *S. pneumoniae* TIGR4 at an MOI of 40. PMNs were placed in the top of the transwell and incubated at 37°C at 5% CO2 for 1.5 hours. PMNs migrated to the bottom of each transwell were enumerated. Technical replicates are pooled from 4 separate experiments (n=4 mice). Asterix indicates significance as determined by unpaired Student’s t-test.

### *S. pneumoniae* infection alters expression of extracellular adenosine pathway components on PMNs

To test if the extracellular adenosine (EAD) pathway regulates the age-driven changes in PMN pulmonary influx following *S. pneumoniae* infection, we first needed to understand its role in young hosts. To do so, we tracked the expression of the different pathway components over time following infection (gating strategy Fig S2). As this is a pulmonary infection and the damage response is initiated in the lungs, we first tracked expression of the EAD producing and degrading enzymes in the lungs of young mice. We measured geometric mean florescence intensity (MFI) on total lung cells as an indicator of expression levels and found that the EAD producing enzymes CD39 and CD73 as well the EAD degrading enzyme ADA were expressed in the lung tissues and that their levels were maintained constant over 18 hours of infection (**Fig S3)**. We had challenges measuring EAD levels in the bronchioalveolar lavage fluid (not shown), so instead we assessed circulating levels and found overall levels of adenosine in the circulation did not significantly change over the course of infection in young mice (**Fig 3**). We then tracked the expression of the four adenosine receptors on the surface of PMNs in the lungs using flow cytometry. We found that at baseline, the majority of PMNs in the lungs of young mice express A1, A2A, and A2B while only about half of PMNs express A3 on the cell surface (**Fig 4A**). To determine if the level of receptor expression on PMNs changes over time, we stained for adenosine receptor expression following infection. We found that all four adenosine receptors showed a significant decrease in expression within the first 18 hours of infection, followed by a recovery of levels comparable to baseline by 48 hpi in young mice (**Fig 4B**). When we measured fold change from baseline, we observed around 2-fold decrease by 6 hours for A2A and A2B and by 18 hours for A1 and A3 followed by recovery for all four receptors by 48 hpi in young mice (**Fig 4C**).

**Figure 3:**
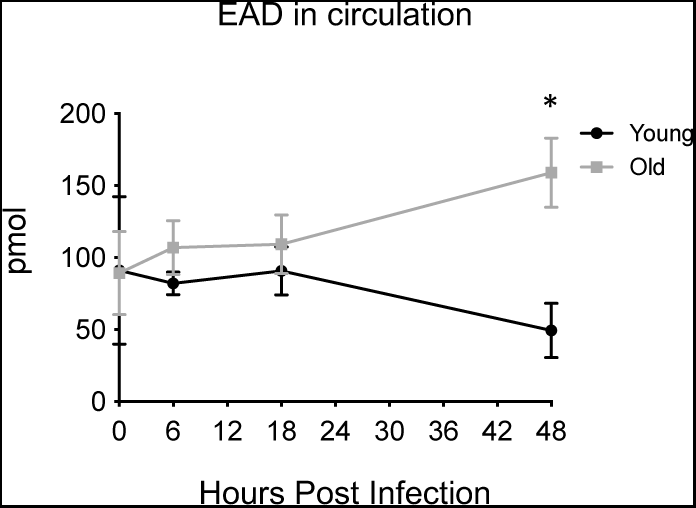
Adenosine levels in circulation change with age over time. Young and old male C57BL/6 mice were infected i.t. with 5×10^4^ CFU *S. pneumoniae* TIGR4 and sera was collected at 6, 18, and 48 hpi. Adenosine levels in the circulation were measured from two separate experiments with n=3 mice at each individual time point. Asterix indicates significantly different from young at the indicated timepoint as calculated by Kruskal Wallis followed by Dunn’s multiple comparison test.

**Figure 4:**
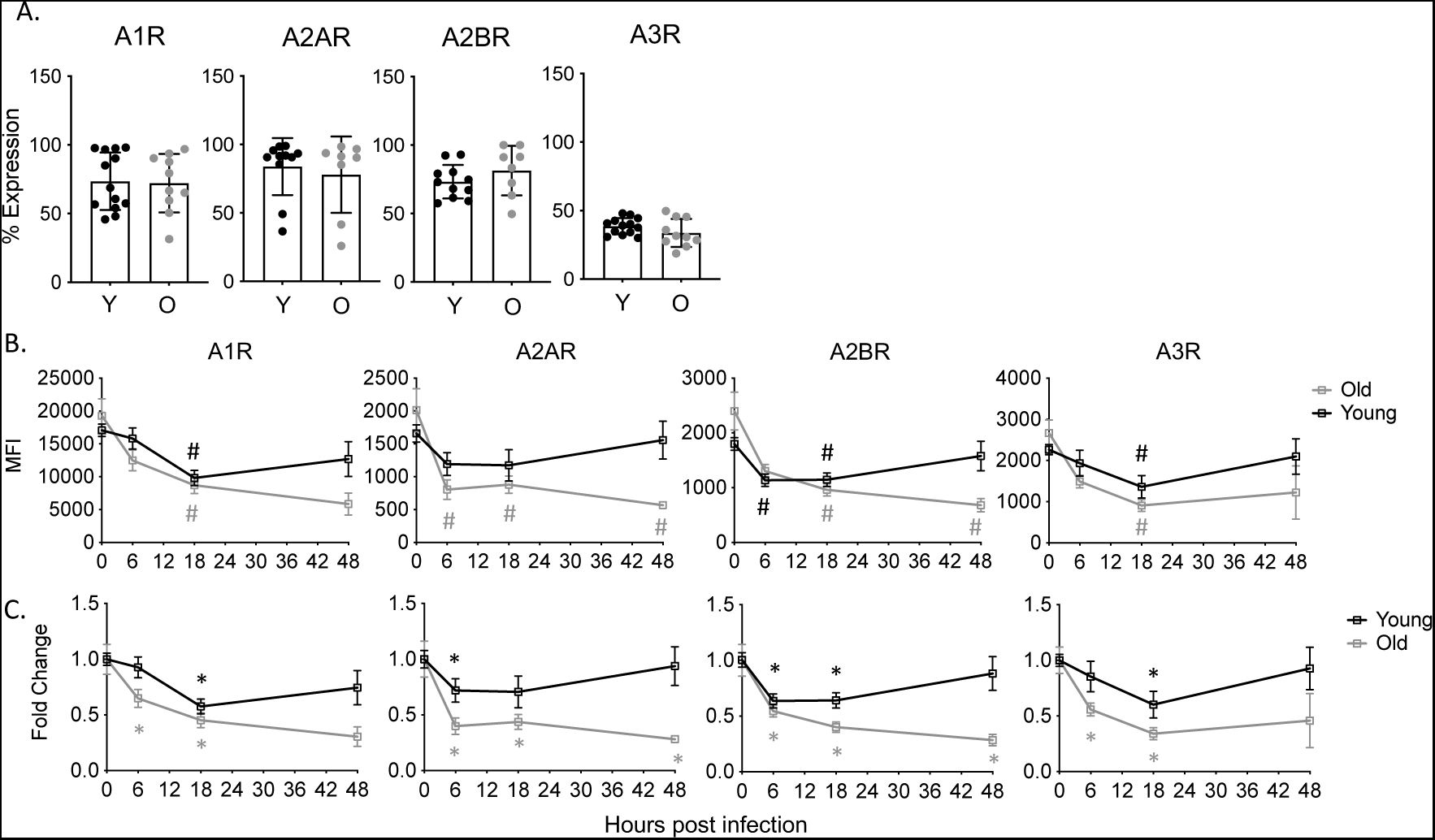
EAD pathway expression on pulmonary PMNs. Young and old male C57BL/6 mice were infected i.t. with 5×10^4^ CFU *S. pneumoniae* TIGR4. The lungs were harvested, and lung digests were stained for PMNs and each adenosine receptor. Percent expression at baseline (A) and MFI over time (B) of the indicated adenosine receptor on PMNs was determined. Replicates are pooled from 6 separate experiments with n=6-12 mice per group. (B) # indicate significance from uninfected baseline as determined by Kruskal Wallis followed by Dunn’s Multiple comparison test. (C) Asterix indicate significant difference from one as determined by a 1-sample t-test.

As PMNs are recruited from the blood, we also measured receptor expression in the circulation. In circulating PMNs of uninfected young mice, A1, A2A, and A2B receptors were highly expressed, however, very few PMNs expressed A3 (**Fig 5A**). When we measured how receptor expression changes over time in circulating PMNs, we found that overall receptor expression remains constant in young mice (**Fig 5B**). Unlike in the lungs, circulating PMNs in young mice expressed A2A, A2B, and A3 at similar level over the course of infection (**Fig 5C**). The expression of A1 in the circulation showed a slight, but significant 1.5-fold decrease at 6 hpi but returned to baseline levels by 18 hpi) (**Fig 5C**). These data show that PMNs in the lungs and circulation of young mice highly express adenosine receptors A1, A2A, and A2B on their surface, and that there are dynamic changes in adenosine receptor expression on PMNs in response to pneumococcal infection.

**Figure 5:**
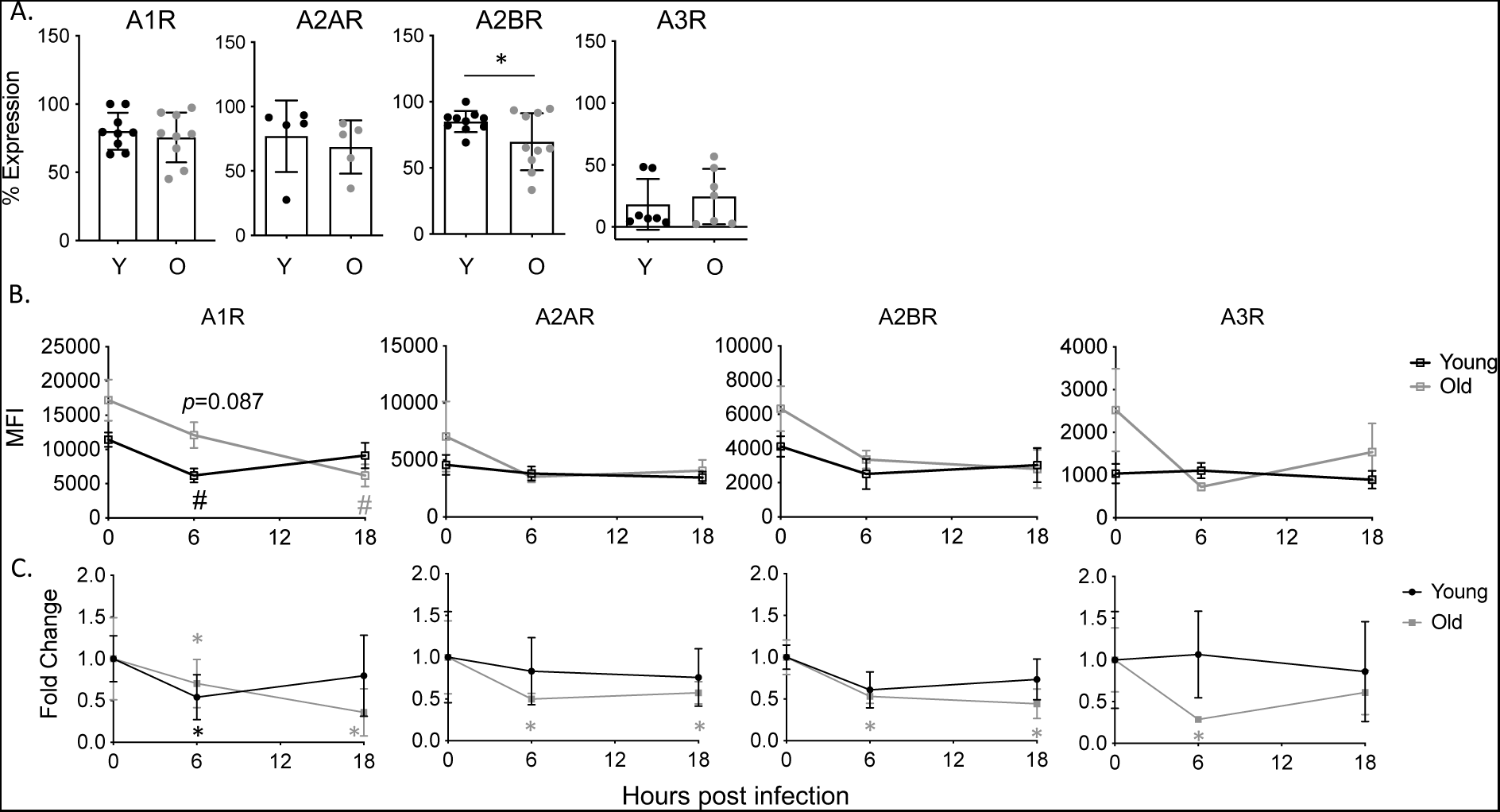
EAD pathway expression on circulating PMNs. Young and old male C57BL/6 mice were infected i.t. with 5×10^4^ CFU *S. pneumoniae* TIGR4. Blood was collected and stained for PMNs and adenosine receptors. Percent expression at baseline (A) and MFI over time (B) was determined using flow cytometry. Replicates are pooled from 4 separate experiments with n=6-9 mice per group. Asterix indicate significant difference as determined by unpaired students t-test (A) and significantly different from one as determined by 1-sample t-test (C). (B) # / p-values indicate significance from uninfected baseline as determined by Kruskal Wallis followed by Dunn’s Multiple comparison test.

### Age-driven changes in EAD-pathway components following *S. pneumoniae* infection

We next wanted to understand how the adenosine pathway components change with host aging. We measured adenosine levels in the circulation of aged mice to see if the dysregulation in PMN influx was due to a lack of extracellular adenosine production. We found that early in infection, there was no significant difference in the amount of adenosine in the circulation when compared to young mice (**Fig 3**). By 48 hpi, aged mice had significantly higher levels of adenosine (**Fig 3**), suggesting exacerbated damage as the infection progressed in aged hosts, which fits with the higher bacterial loads observed in aged hosts (**Fig 1**). When we measured adenosine pathway enzymes in the lungs, we found that at baseline, the lungs of aged mice express significantly higher amounts of the EAD producing enzymes CD73 and CD39, while the levels of the EAD degrading enzyme ADA were higher, but did not reach statistical significance (**Fig S3**). When we tracked enzyme expression through the course of infection, we found that unlike young controls, the expression of all three enzymes had significantly decreased by 18 hpi in lungs of aged mice (**Fig S3**).

Next, we analyzed receptor expression on PMNs in the lungs and circulation. We found that in the lungs and blood at baseline there was no change in the percentage of PMNs expressing A1, A2A, A2B, or A3 when compared to young mice with the majority of cells expressing A1, A2A, and A2B (**Fig 4A** and **Fig 5A**). To look at receptor expression changes overtime, we measured MFI and calculated fold change from baseline. In both lung (**Fig 4B**) and blood (**Fig 5B**) there was no significant difference in receptor expression between young and aged mice and the expression of A1 receptor trended to be higher on circulating PMNs at 6 hpi in aged mice compared to young controls (**Fig 5B**). We did find that PMNs in the lungs of aged mice decreased levels of adenosine receptors early in infection respective to their own baseline, however, unlike in young mice where receptor expression returned to baseline levels by 48 hours, with age there was no rebound of adenosine receptor expression at this later time point (**Fig 4C**). A similar pattern of decreased receptor expression over time in aged mice on circulating PMNs was also seen, where circulating PMNs maintained a 2-fold reduction in expression of A1, A2A and A2B following infection (**Fig 5C**). These data show that there is no defect with age in production of extracellular adenosine and or adenosine receptor expression on PMNs. Rather, the decrease in adenosine receptor expression on PMNs during the course of infection is more pronounced and may result in decrease responsiveness to adenosine signaling in the aged hosts.

### A2B adenosine receptor plays no role in PMN pulmonary influx following *S. pneumoniae* **infection**

To determine if the individual adenosine receptors played a role in PMN pulmonary migration, we focused on the receptors highly expressed on PMNs (A1, A2A and A2B). We first examined the A2B receptor. To examine the role of this receptor in PMN influx to the lungs *in vivo,* WT C57BL/6 and A2BR^-/-^ mice were infected intratracheally with 2.5×10^4^ CFU *S. pneumoniae* and the total number of PMNs in the lungs was determined by flow cytometry at 6, 18, and 48 hpi. We found that there was no difference in the total amount of PMNs in the lungs at any of the time points tested between the mouse strains (**Fig 6**). Bacterial burden in the lungs also did not differ (data not shown and supporting what was previously published (38)) and even when normalized for bacterial numbers, PMN influx into the lungs in response to *S. pneumoniae* infection was comparable between WT and A2BR^-/-^ mice (**Fig S1B**). These data suggest that A2B receptor signaling does not regulate PMN influx to the lung of *S. pneumoniae* infected mice.

**Figure 6:**
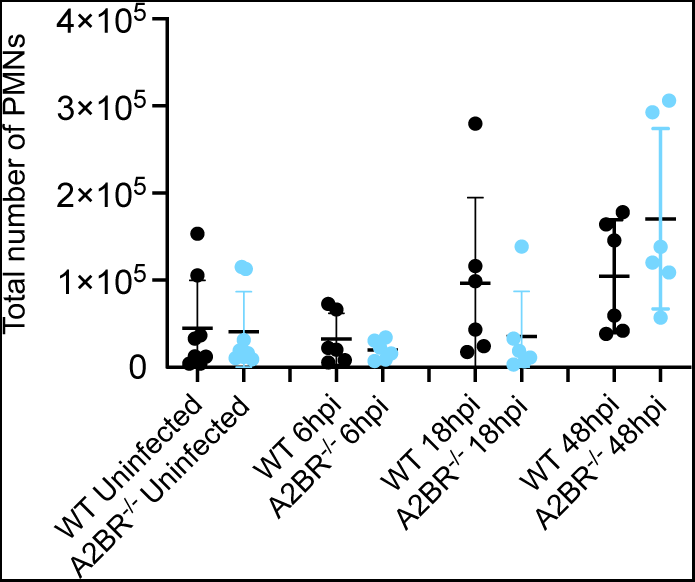
A2B receptor does not affect PMN pulmonary numbers *in vivo.* WT and A2BR^-/-^ mice were infected i.t with 5×10^4^ CFU *S. pneumoniae* TIGR4. At the indicated timepoints following bacterial challenge, lungs were harvested, digested, and stained for PMNs and analyzed by flow cytometry. Data are pooled from 4 separate experiments with n=6-8 mice per group.

### A2A adenosine receptor plays no role in PMN pulmonary influx following *S. pneumoniae* **infection**

Next, we examined the role of A2A receptor signaling in PMN influx to the lungs following infection. To examine the role of this receptor *in vivo*, A2AR^-/-^ and WT BALB/c controls were infected intratracheally with 2.5×10^4^ CFU *S. pneumoniae* TIGR4 and total number of PMNs in the lungs was determined at 6 and 48 hpi by flow cytometry. There was no difference in the number of PMNs in the lungs of A2AR^-/-^ or WT mice at either time point (**Fig 7A**). To confirm that this result was not due to differences in bacterial burden, we analyzed *S. pneumoniae* CFU in both the lung (**Fig 7B**) and blood (**Fig 7C**) and found no difference in bacterial burden across mouse strains. Additionally, there was no difference in survival following infection between A2AR^-/-^ and WT mice (**Fig 7D**). Since these experiments were done using mice on a Balb/c background, we also analyzed the role of A2A in C57BL/6 mice using a specific pharmacological A2A receptor inhibitor, 3,7-Dimethyl-1-propargylxanthine. We first tested the role of A2A early in infection by treating mice with the A2A receptor antagonist or vehicle control 18 hours prior to bacterial challenge (PRE). We found no difference in total number of PMNs in the lungs (**Fig S4A**), total bacterial burden in the lung (**Fig S4B)** or bacteremia (**Fig S4C)** by 48 hpi. Similarly, inhibition of A2A prior to infection had no effect on mouse survival (**Fig S4D).** As A2A is a lower affinity receptor that may play a role later in infection as damage and EAD levels build up, we also tested its role later in infection, by timed inhibition 18 hours post (POST) bacterial challenge. Again, we found no difference in total bacterial burden in the lung (**Fig S4B)** or bacteremia (**Fig S4C)** by 48 hpi in POST treated mice. However, there was a decrease in survival in the mice treated with A2A receptor inhibitor post infection (**Fig S4D**) suggesting that A2A receptor may play a protective role later in infection. Overall, these data suggest that A2A receptor signaling does not regulate PMN influx to the lungs of young mice following *S. pneumoniae* challenge.

**Figure 7:**
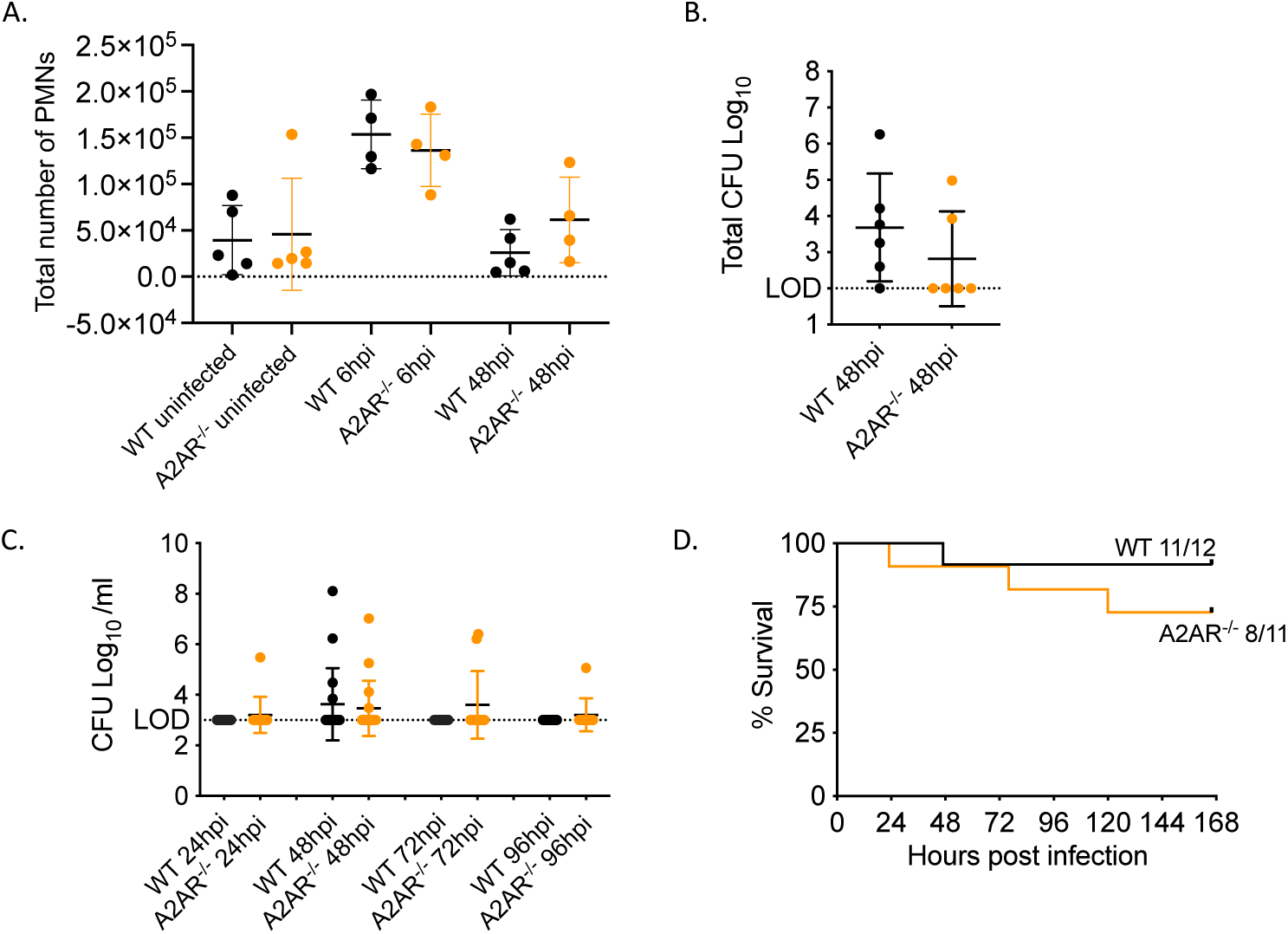
A2A receptor does not affect PMN pulmonary numbers or host resistance to infection. (A) WT and A2AR^-/-^ mice were infected i.t with 5×10^4^ CFU *S. pneumoniae* TIGR4. At the indicated timepoints following bacterial challenge, lungs were harvested, digested, and stained for PMNs and analyzed by flow cytometry. Data are pooled from 3 separate experiments with n=4-6 mice per group. Lungs (B) and Blood (C) were harvested at indicated timepoints and plated on blood agar plates and CFU was enumerated. Data were pooled from 2 separate experiments with (C) n=11-12 mice per group and (D) n=6 mice per group. (E) WT and A2AR^-/-^ mice were infected as in (A) and monitored for survival. Data are pooled from 2 experiments with n=11-12 mice per group. No significant differences were found in survival as measured by Log-Rank (Mantle Cox) test.

### A1 receptor signaling is required for PMN pulmonary influx in response to *S. pneumoniae* infection

To determine the role of A1 receptor signaling in PMN migration, young mice were treated i.p. with a specific A1 receptor inhibitor (whose specificity we confirmed in prior publications (28, 33)) or vehicle control. Mice were then infected intratracheally with 2.5×10^4^ CFU *S. pneumoniae* TIGR4 one day post drug treatment. Total PMN influx to the lungs of infected mice was measured by flow cytometry at 6, 18, and 48 hours. We found that inhibition of A1 resulted in a significant delay in PMN influx to the lungs at 6 hpi (**Fig 8A-B**). At 6 hpi, there was a significant decrease in total PMN numbers in the lungs (**Fig 8A**) and the amount of MPO (a marker of PMNs) in lung homogenates (**Fig 8B**) of A1 inhibited mice. To understand the mechanism of the decline in PMN recruitment, we examined the expression of CD18 and CXCR2 on the surface of PMNs in the circulation. We found no difference in the numbers of circulating PMNs in A1 receptor inhibited versus vehicle treated mice (**Fig S5)**. We further found that while A1 receptor inhibition had no effect of CD18 expression (**Fig 8C**), it significantly reduced CXCR2 expression at 6 hpi (**Fig 8D**), the time point at which PMN number in the lungs is lower (**Fig 8A**). As CXCR2 was reported to be required for trans-endothelial migration in response to pneumococcal infection (11, 14, 39–41), we directly assessed the role of A1 on PMN trans-endothelial migration. Young mice were treated i.p. with an A1 receptor inhibitor, and PMNs were then isolated, and trans-endothelial migration was measured using the same *in vitro* transwell assay described above. A1 receptor inhibited PMNs displayed a significant decrease in their ability to migrate trans-endothelially compared to VC treated PMNs, indicating that A1 signaling on PMNs is required for efficient trans-endothelial migration (**Fig 8E**). These data suggest that A1 adenosine receptor signaling is required for recruitment of PMNs to the lungs in response to *S. pneumoniae* infection.

**Figure 8:**
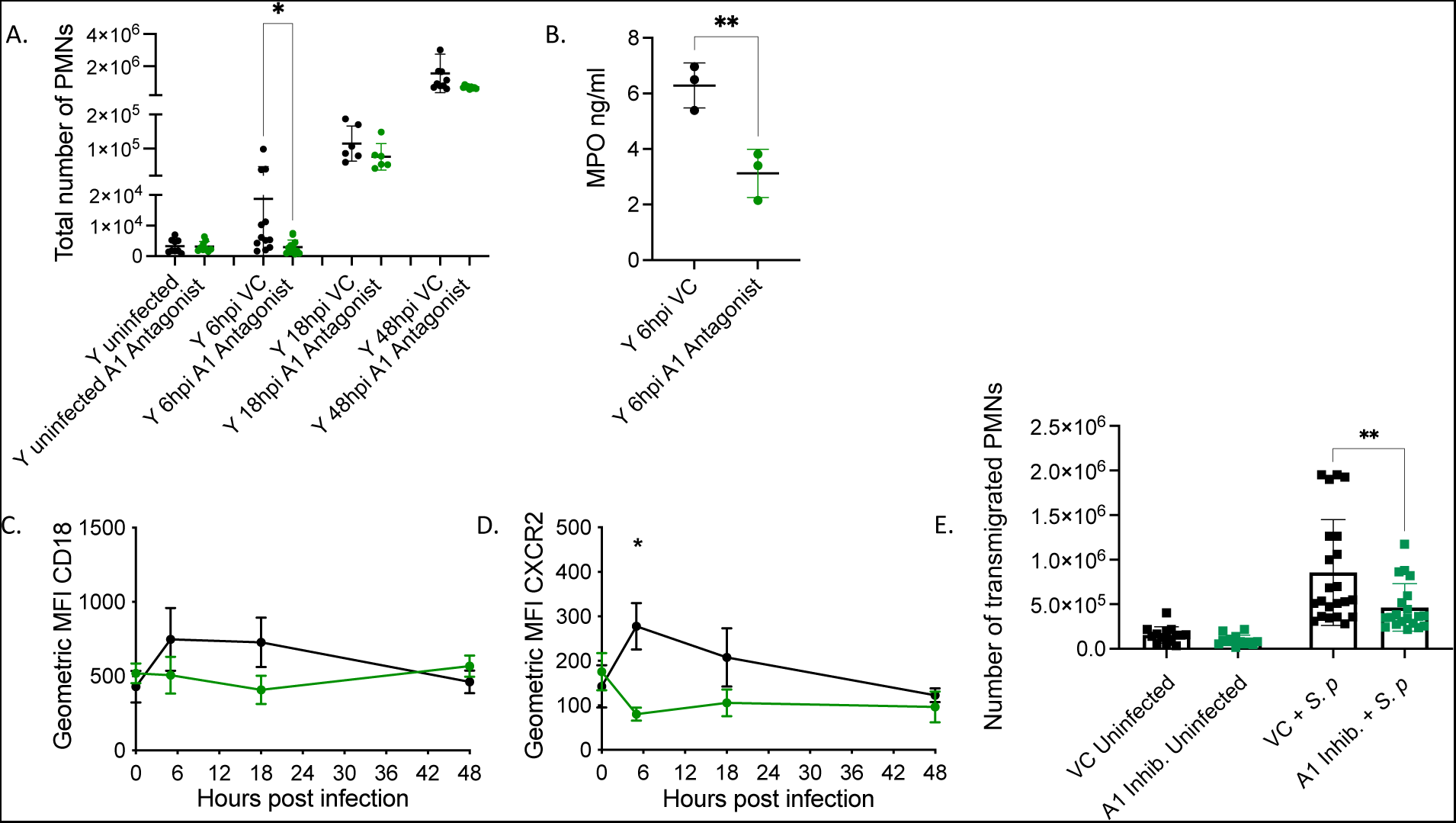
A1 receptor signaling is required for PMN pulmonary influx. (A) Young, male mice were treated i.p with A1 inhibitor DPCPX and infected i.t with 5×10^4^ CFU *S. pneumoniae* TIGR4. At the indicated time points, lungs were harvested, digested, and stained for PMNs (CD45+, Ly6G+ CD11b+) (A) and analyzed by flow cytometry. (B) Lungs of a separate set of treated and infected mice were harvested 6 hpi, homogenized and MPO levels were determined by ELISA. (C-D) Blood was harvested from infected and A1 treated mice (same mice as A) at the indicated time points and stained for PMNs and CD18 (C) or CXCR2 (D). (A, C and D) Data are pooled from 3 separate experiments with n=6-12 mice per group. (A and D) Asterix indicate significance as determined by Kruskal Wallis followed by Dunn’s multiple comparison test. (B) Asterix indicate significance as determined by unpaired students t-test. (E) Young, male C57/B6 mice were treated i.p. with A1 inhibitor DPCPX or vehicle control (VC) and PMNs were isolated from the bone marrow. PMN migration across primary endothelial cells in response to *S. pneumoniae* TIGR4 infection was measured. Technical replicates are pooled from 4 separate experiments. Asterix indicates significance as determined by Mann-Whitney test.

### Activation of A1 receptor signaling rescues the ability of PMNs from old mice to be recruited to the lungs following *S. pneumoniae* infection

To better understand if adenosine receptor signaling can be targeted to reverse the age associated dysregulation in PMN pulmonary influx following pneumococcal infection, aged mice were treated with specific adenosine receptor agonists to stimulate receptor signaling and then infected intratracheally with 2.5×10^4^ CFU *S. pneumoniae* TIGR4. We first tested A2A receptor as it had a role in controlling survival of young hosts later in infection. We found that activating A2A in aged mice at 18 hpi had no effect on the total number of PMNs in the lungs (**Sup. Fig 6A**), or the number of bacteria in the lungs (**Sup. Fig 6B**), or blood (**Sup. Fig 6C**) by 48 hours post challenge.

We then focused on the role of A1 as it was required for early influx of PMNs in young hosts. We tested the effect of A1 agonism in aged mice and determined the total number of PMNs in the lungs at 6 hpi by flow cytometry. We found that agonism of the A1 receptor significantly increased the total number of PMNs in the lungs of aged mice 6 hpi when compared to uninfected mice (**Fig 9A**). Additionally, at 6 hpi, a separate set of A1 agonist treated mice had significantly higher levels of MPO in the lungs compared to vehicle control treated mice (**Fig 9B**). Importantly, A1 receptor agonism decreases bacterial burden in the lungs of aged mice compared to untreated controls (**Fig 9C**). Taken together, these data demonstrate that A1 adenosine receptor signaling can be targeted to reverse the age-related decrease in PMN pulmonary influx and reduce bacterial burden in the lungs of infected aged mice.

**Figure 9:**
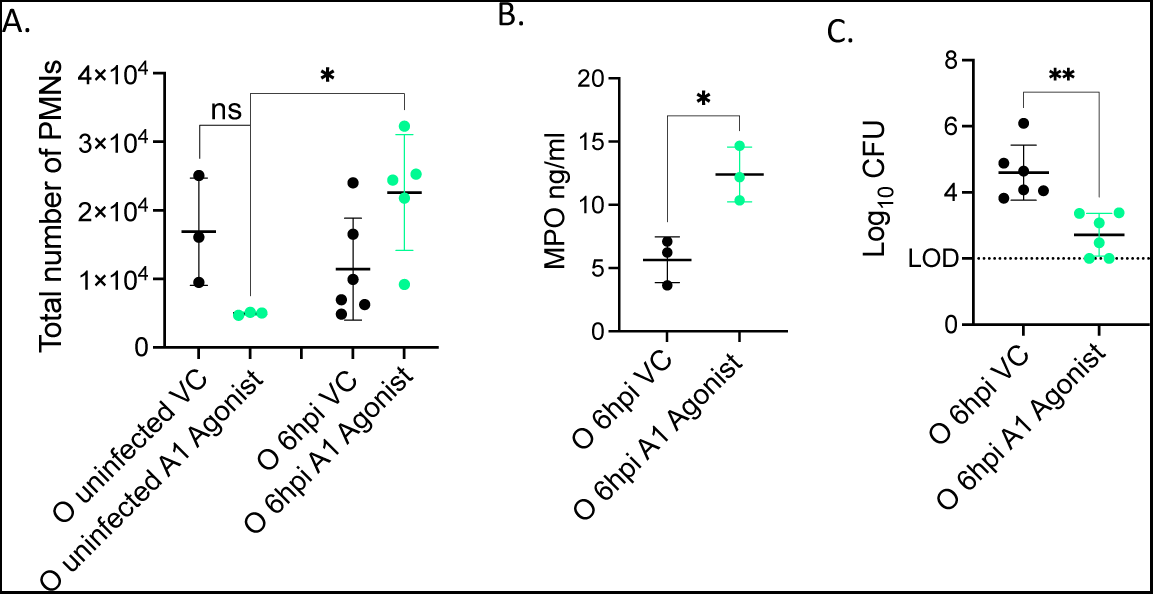
A1 receptor signaling rescues defect in early PMN pulmonary influx in aged mice. (A) Old male mice were treated i.p. with A1 agonist 2-Chloro-N6-cyclopentyladenosine or Vehicle Control and infected i.t. with 5×10^4^ CFU *S. pneumoniae* TIGR4 for 6 hours. Lungs were harvested, digested, and stained for PMNs and analyzed by flow cytometry. Asterix indicates significance as determined by One-way ANOVA followed by Sidak’s multiple comparison test. ns indicates non-significant. (C) Lung homogenates from infected mice were plated on Blood agar and bacterial CFU was enumerated. Data are pooled from two separate experiments and with n=3 uninfected and n=6 infected mice per group. (B) Lungs of a separate set of treated and infected mice were harvested 6 hpi, homogenized and MPO levels were determined by ELISA. Data are pooled from n=3 mice. (B and C) Asterix indicates significance as determined by unpaired Student’s t-test.

## DISCUSSION

Cellular damage resulting from infection is proposed to shape the outcome of host-pathogen interactions (42–44) and damage-associated molecular patters (DAMPS) can act as signals that initiate the immune response (45). One such molecule is extracellular adenosine (EAD). At baseline homeostatic levels, adenosine levels in extracellular microenvironments are very low, however, upon tissue damage resulting from insults such as infection, EAD levels can rise more than 10 fold (18). We previously found that EAD production and signaling are crucial for host resistance against pneumococcal infection (10, 20, 21, 38, 46). We show here that EAD signaling through the high affinity A1 receptor is required for initiating the recruitment of PMNs into the lungs following bacterial challenge. This initiation of early PMN recruitment was defective in aged hosts but could be reversed by activation of A1 receptor signaling, suggesting that with age there are changes in extracellular adenosine signaling that impair the ability of PMNs to rapidly respond to pneumococcal infection. One of the hallmarks of aging is altered intercellular communication, including endocrine and neuronal signaling (7) and the findings here suggest that damage signaling is also altered with aging and contributes to the age-driven changes in immune response to infection.

PMNs have a dynamic role during pneumococcal pneumonia with their early presence in the lungs being protective vs later persistence being detrimental (47). In murine models of infection, PMN influx to the lungs within the first 12 hpi correlated with control of bacterial burden and PMN depletion prior to infection resulted in increased bacterial burden in the lung and host lethality (48, 49). In humans, patients with neutropenia are at an increased risk for pneumonia (13). These findings indicate that early influx of PMNs to the lungs is required for host protection against pneumococcal pneumonia. In contrast, PMN persistence in the pulmonary environment promotes disease (10, 15, 17). Depletion of PMNs 18 hours after pneumococcal lung infection resulted in reduced bacterial numbers and reduced lethality (10). These findings demonstrate that the ability of PMNs to kill pneumococci is altered during infection (10). Here we found that with aging there is a delay in PMN influx early within the first 6 hours of infection accompanied by persistent presence in higher levels later in infection compared to young mice infected with the same dose of *S. pneumoniae*. This delay left aged mice more susceptible to disease as it was accompanied by increased bacterial burden in the lungs throughout the course of infection and higher incidence of bacteremia compared to young mice. Therefore, with age there is a delay in the protective early PMN recruitment to the site of infection. The majority of studies examining PMN influx in response to infection in aged hosts have focused on later timepoints (9, 10, 17, 50, 51). In humans, elderly pneumococcal pneumonia patients had higher percentages of neutrophils in terminal lung biopsies compared to younger patients (16) and in several animal models of bacterial and viral pneumonia, PMN numbers in the lungs of aged mice were higher than in young controls at later timepoints (9, 10, 17, 50, 51), similar to what we found here. The dynamics of earlier influx and how that is altered with aging have not been as thoroughly addressed in the literature and there are no data from human pneumococcal pneumonia patients. Our findings suggest that the kinetics of PMN influx to the lungs in response to pneumococcal pneumonia is altered with age.

Following lung challenge, *S. pneumoniae* localizes within the airways and binds to lung tissues as well (10, 17). For PMNs to reach the site of infection in the lungs, they have to move from the circulation across the endothelium, enter the interstitial space, then migrate across the lung epithelium into the airways (11, 52). We found here that PMN migration across pulmonary endothelial cells in impaired with aging. Prior work found that PMNs from elderly human donors did not show defects in transepithelial migration in response to pneumococcal infection (53) but rather that chemotaxis (or directional movement) of PMNs is diminished with aging (51, 54–58). PMNs isolated from both healthy controls and pneumonia patients above 60 years of age displayed diminished chemotaxis compared with younger individuals (54, 55) which was linked to over activation of phosphoinositide-3-kinase (PI3K) signaling (54). PI3K is controlled downstream of GPCRs including adenosine receptors (54, 59). Here we found that A1 receptor signaling on PMNs is required for their migration across pulmonary endothelial cells in response to infection and that activating A1 receptor reverses the defect in early PMN pulmonary influx observed in aged hosts. These findings are supported by prior work showing that A1 receptor signaling on PMNs is required for their adherence to the endothelium in response to N-Formylmethionyl-leucyl-phenylalanine (fMLP) or Phorbol 12-myristate 13-acetate (PMA) stimulation (60, 61). In this study we found that A1 receptor signaling was needed for the ability of PMNs to upregulate the expression of the chemokine receptor CXCR2 early on in response to infection. During *S. pneumoniae* infection, CXCR2 is required for PMN influx to the lungs (39) and in aged mice, expression of CXCR2 on PMNs was reported to be lower than young hosts (56). Therefore, it is possible that activating A1 receptor signaling in aged hosts boosts PMN migration by upregulating expression of CXCR2.

The role of A1 receptor signaling in PMN recruitment to the lungs seem to play opposing roles during pathogen driven vs sterile injury. In LPS or ischemia reperfusion lung injury, A1 receptor signaling attenuated PMN trafficking to the lungs (62–65). In contrast and similar to what we observed here, A1 receptor signaling was required for PMN pulmonary influx during infection with H1N1 influenza virus (66). This highlights that pathogens can alter the dynamics of the immune response and that the role of the adenosine signaling may differ in sterile vs. infection driven injury.

Extracellular adenosine is known to regulate PMN function (18) and play an important role in host resistance against pulmonary infections (67). Aging is associated with aberrant PMN responses and overall increased susceptibility to pulmonary infections (47). However, surprisingly, the role of EAD in immunosenescence remains largely unexplored. In prior work we found that there are age driven changes in the expression of the enzymes that make and break EAD on PMNs and that EAD signaling on PMNs could be targeted to reverse the decline in their antibacterial activity (33). In this study, to examine the role extracellular adenosine signaling in the age-related delay in PMN influx we tracked how this pathway changes with host aging. We found that circulating levels of adenosine were no different in young and aged mice, in fact aged mice had significantly more adenosine in circulation at 48 hpi. While this increase in adenosine later on in infection may reflect the increased damage that is occurring in these mice (due to higher bacterial burden), these data also indicate that it is not that adenosine is not being produced with age, but that EAD signaling in PMNs of aged mice may be dysregulated. This is in line with prior work showing that aging is associated with defects in overall GPCR signaling and that alterations in EAD receptor signaling contribute to the age-associated decline of brain and heart organ functions (25) (68) (23). When looking at adenosine receptor expression on the surface of PMNs from aged mice following pneumococcal infection, there was a decrease in receptor expression from baseline in both you and aged mice. However, in PMNs isolated from young mice these receptor levels rebounded by 48hpi, while receptor expression on PMNs from aged mice did not recover. Following activation by an agonist, GPCR signaling is desensitized by internalization of the receptor into the cell, this is a result of interaction of the G protein with β-arrestins to stop signaling and additional scaffold proteins which guide trafficking the internalized GPCR (69–71). The decrease in adenosine receptor expression following pneumococcal infection may be due to this process of GPCR desensitization. Following desensitization, GPCRs may be degraded but may also recycle back to the cell membrane surface for resensitization, where the receptor can be activated again (70). In PMNs isolated from aged mice adenosine receptor expression does not rebound following the initial drop post infection. This would indicate that EAD receptors are not being recycled back to the cell surface, and throughout infection, PMNs of aged mice may be less responsive to activation by adenosine. This could be in part be driven by the consistently elevated levels of EAD later in infection observed in old but not young hosts. Additionally, the differences in receptor levels between young and aged mice later in infection could indicate that there are changes in GPCR desensitization and re-sensitization in PMNs with host age.

A balanced PMN response where PMNs are recruited to the lungs and resolve at the proper time is essential for effective clearance of pneumococcus and efficient host protection (12, 13, 15, 16). In prior work, we found that in young mice, EAD production by CD73 was needed for regulation of PMN influx later on during infection, where by day 2 following pulmonary challenge, CD73^-/-^ mice had significantly more PMNs in the lungs compared to wildtype controls (10). We hypothesized that extracellular adenosine signaling via the low affinity receptors A2A and A2B receptors regulate PMN resolution from the lungs later in infection when adenosine levels are higher due to increased inflammation and host damage. It has been shown previously that under hypoxia conditions adenosine produced by the endothelium reduced PMN accumulation by activating A2A and A2B receptors on the PMN surface (72). Surprisingly, we found here that young mice lacking A2A or A2B receptors showed no changes in PMN presence in the lungs thorough the course of pneumococcal infection. As both A2A or A2B receptors are expressed on PMNs and are coupled to Gs (18), it is possible that they play a redundant in PMN movement and that removal of both would be needed to observe a significant effect. An alternative explanation is that the levels of adenosine receptors do decline on PMNs through infection even in young hosts, suggesting that PMNs may become less responsive to adenosine signaling with time.

In summary, we have shown that A1 receptor signaling is required for early PMN influx to the lungs following pneumococcal infection. Importantly, activation of the A1 receptor using A1 receptor agonist, 2-Chloro-N6-cyclopentyladenosine increased PMN influx to the lung in aged mice, which resulted in lower bacterial burden in the lung. This enhanced ability to control bacterial infection could be a combination of increase PMN influx as shown here and improved function as previously reported (33). These findings suggest that extracellular adenosine signaling may be dysregulated with host aging and that pharmacologically targeting this pathway can enhance the immune response in aged mice leading to more effective clearance of pneumococcus. Identifying a potential target to enhance the immune response to pneumococcal pneumonia in aged hosts is important as this disease persists in older adults despite the availability of antibiotics and vaccines (3, 5, 6). In addition to being a potential pharmacological target, the extracellular adenosine pathway is an important pathway to understand in the context of host-pathogen interaction as it could be a marker of host damage. This study adds dysregulation of the EAD pathway with age under the aging hallmark of altered intercellular signaling, which in turn controls immunosenescence.

## DECLARATIONS

### Ethics approval and consent to participate

All animal experiments were conducted in accordance with the Institutional Animal Care and Use Committee guidelines under the approved protocol number MIC33018Y

### Consent for publication

Not applicable

### Availability of data and materials

All data generated or analyzed during this study are included in this published article [and its supplementary information files].

### Authors’ contributions

SRS designed research, conducted research, analyzed data, and wrote paper. SES designed research, conducted research, and analyzed data. EYIT, APL and MB conducted research and analyzed data. ENBG designed research, wrote the paper, and had responsibility for final content. All authors read and approved the final manuscript.

## Acknowledgements

We would like to thank Dr. Michael Battaglia for critical feedback on the manuscript.

