## Supplemental Materials for "Activating A1 adenosine receptor signaling boosts early pulmonary neutrophil recruitment in aged mice in response to *Streptococcus pneumoniae* infection"

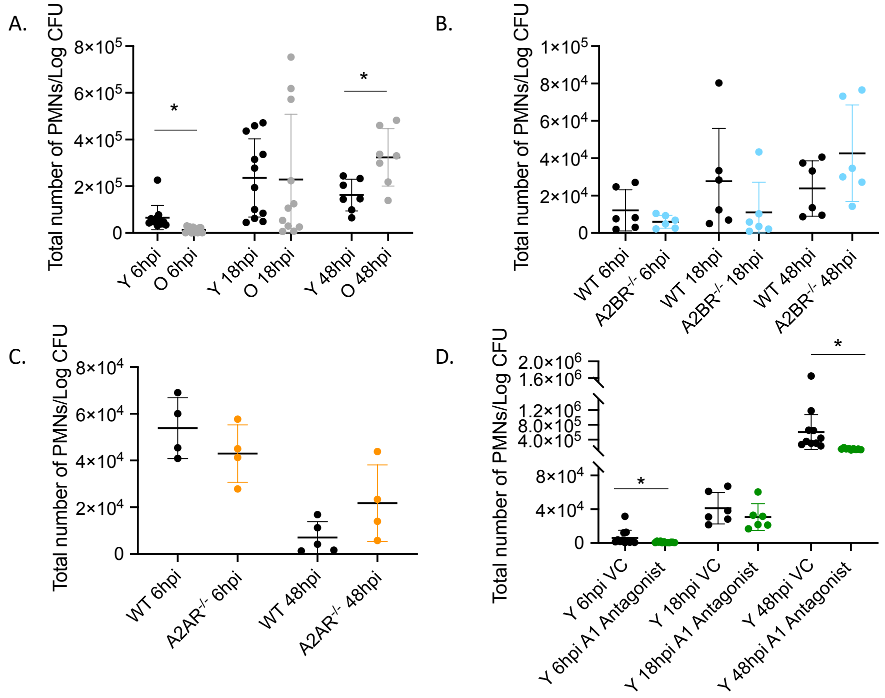


**Supplemental Figure 1: Total PMNs/Log CFU in Lung.** (A)Young and old male C57BL/6 mice (B), A2BR$^{-/-}$ mice, (C) A2AR$^{-/-}$ mice and (D) Young C57BL/6 mice treated with A1 inhibitor were infected intra-tracheally (i.t.) *S. pneumoniae* TIGR4. Lungs were harvested, digested and stained for PMNs and analyzed by flow cytometry at the indicated timepoints following infection. Lung samples were plated on blood agar and CFU was enumerated at the indicated timepoint post infection. (A-D) For each mouse, the number of PMNs in the lungs were divided by the total CFU per lung. (A) Data are pooled from 6 separate experiments with a total of n=7 mice at 48-hour time point and n=12 mice at all other time points. (B) Data are pooled from 4 separate experiments with n=6-8 young mice per group. (C) Data are pooled from 2 experiments with n=11-12 young mice per group. (D) Data are pooled from 3 separate experiments with n=6-12 mice per group. Asterix indicates significance as determined by Kruskal Wallis followed by Dunn's Multiple comparison test.

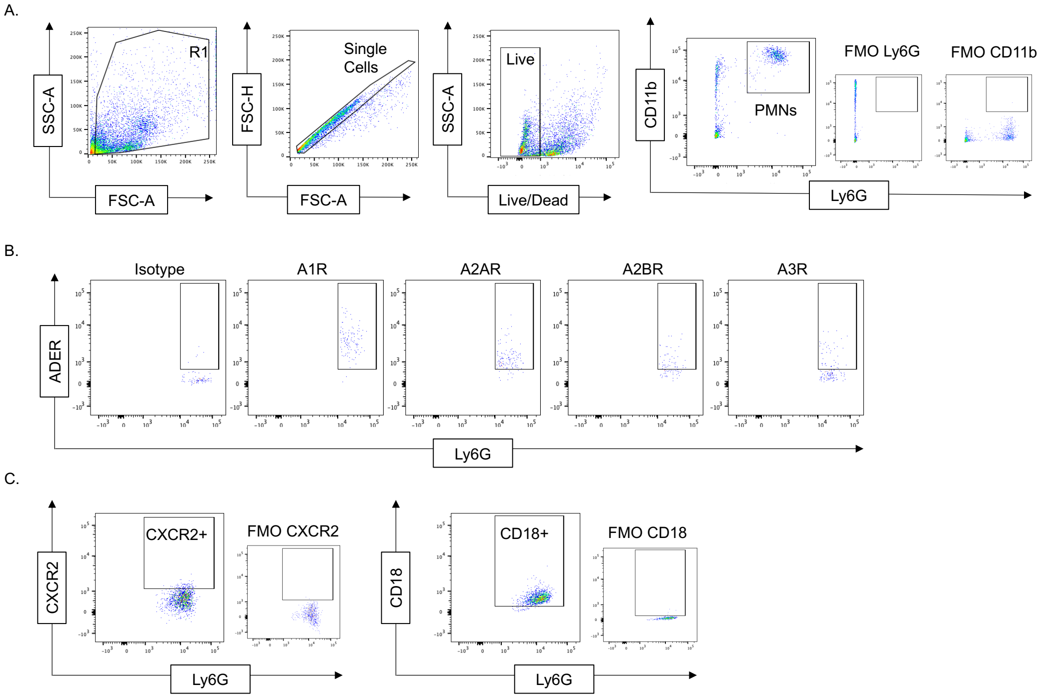


**Supplemental Figure 2: Gating Strategy.** Lungs and blood were harvested, and receptor expression was determined by flow cytometry. Representative gating strategies are shown. (A) single cell, live, PMNs (Ly6G$^+$, CD11b$^+$) were gated on and (B) expression of the different adenosine receptors as well as (C) expression of CXCR2 and CD18 on PMNs was determined. Abbreviations: SSC-A= side scatter peak area, FSC-A= forward scatter peak area, FSC-H= forward scatter peak height, FMO= fluorescence minus one.

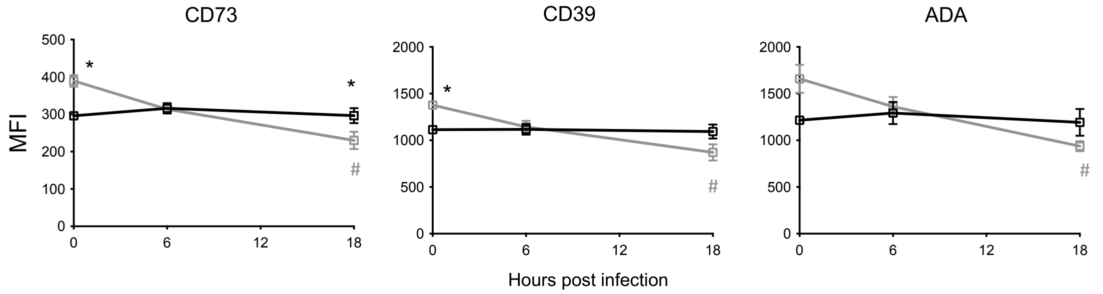


**Supplemental Figure 3: EAD pathway enzyme expression in lungs.** Young and old male C57BL/6 mice were infected i.t. with 5x$10^4$ CFU *S. pneumoniae* TIGR4. The Lungs were harvested, and lung digests were stained for the indicated adenosine producing and degrading enzymes. MFI on the surface of PMNs was determined over time using flow cytometry. Asterix indicates significance between young vs old at the indicated timepoint and # indicates significance for each group from its uninfected baseline as determined by Kruskal Wallis followed by Dunn's multiple comparisons test. Data are pooled from 2 separate experiments with n=12-14 mice per group.

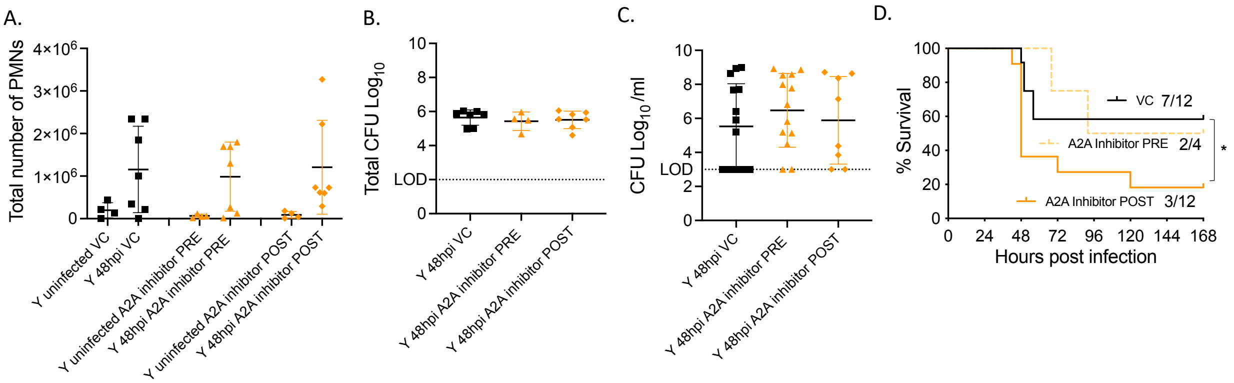


**Supplemental Figure 4: Signaling via A2A receptor does not affect PMN pulmonary numbers or host resistance to infection.** Young, male mice were treated i.p. with specific A2A receptor inhibitor or vehicle control (VC) either 18 hours prior to infection (PRE) or 18 hpi (POST). Mice were then infected i.t. with $5x10^4$ CFU *S. pneumoniae* TIGR4. (A) Lungs were harvested 48 hpi, digested, and stained for PMNs. Samples were analyzed by flow cytometry. (B) Lung homogenates and (C) blood samples were plated on blood agar and bacterial CFU was enumerated. (A-C) Data are pooled from 3 separate experiments with (A) n=4-8 per group, (B) n=4-6 per group, (C) n=8-14 mice per group. (D) Following treatment and infection mice were monitored for survival. Asterix indicates significance as determined by Log-Rank test. Fractions indicate the number of surviving mice over the number of total mice per group. Data are pooled from two separate experiments.

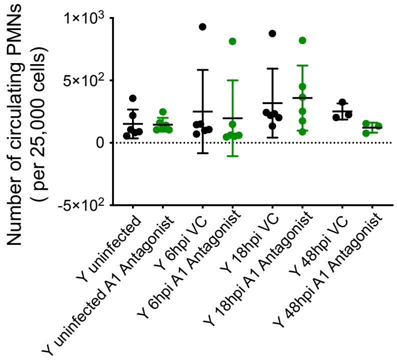


**Supplemental Figure 5: A1 receptor inhibition does not affect the number of circulating PMNs.** Young, male C57/B6 mice were treated i.p. with specific A1 inhibitor and infected i.t with $5x10^4$ CFU *S. pneumoniae* TIGR4. At indicated time points blood was harvested and stained for PMNs (Ly6g+, CD11b+) and analyzed by flow cytometry. Data are pooled from 3 separate experiments with n=6-12 mice per group.

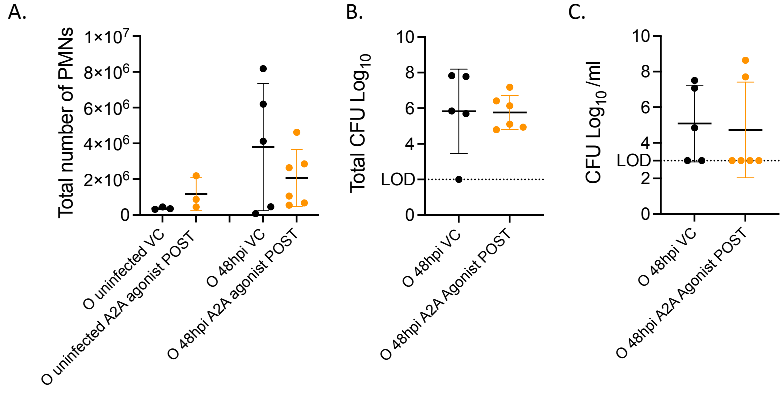


**Supplemental Figure 6: A2A receptor signaling does not affect late PMN pulmonary migration in aged hosts.** Old mice were infected at $2x10^4$ CFU with *S. pneumoniae* TIGR4 and 18 hpi treated i.p. with specific A2A agonist. 48 hpi lungs were harvested and total number of PMNs were determined by flow cytometry (A). Lungs were homogenized (A) and blood was collected (C) and plated on blood agar plates and CFU was enumerated. Data are pooled from n=3-6 mice per group.
